# Structural Evolution of Trimeric Gamma-Type Retroviral Envelope Proteins

**DOI:** 10.1101/2023.10.07.561363

**Authors:** Isidro Hötzel

## Abstract

The envelope glycoproteins (Env) of retroviruses are highly diverse due to their relatively rapid evolution and the ancient origin of retroviruses. Sequence similarity among the Env of retroviruses is observed in the transmembrane subunit (TM) of Env mediating membrane fusion, whereas this similarity is more limited or absent in the surface subunit (SU) mediating receptor binding. Recent structural modeling of the SU of major retroviral groups identified a structurally conserved β-sheet domain in SU and the SU-equivalent of filoviruses, GP1, that is differentially expanded to generate the structural diversity observed in this protein family. Here, trimeric Env structures from three major exogenous and endogenous gamma-type retroviral Env classes were modeled with AlphaFold2 multimer to address the evolution of trimeric Env in the retroviruses. The models show a close structural relationship between the trimeric Env proteins of gammaretroviruses and alpharetroviruses and the trimeric filoviral spike protein, with an extended hydrophobic structure anchoring SU into TM in retroviral Env occluding the intersubunit disulfide bond. In addition, the murine leukemia virus (MLV) Env trimer model indicates that the type-C gammaretroviral receptor binding domains (RBD) are positioned on the side of the Env trimer pointing towards the viral envelope membrane, unlike those of the gammaretroviral RDR interference group, which are positioned closer to the apical end of the Env trimer. The trimeric MLV Env structural model, including RBD position, is consistent with previous trimeric MLV Env electron-density maps. The structural models provide insights into the structure, function and evolution of trimeric retroviral gamma-type Env.

## INTRODUCTION

The envelope glycoprotein (Env) of retroviruses mediates infection by inducing fusion between the viral and host-cell membranes. The Env of retroviruses also plays critical roles in the physiology of reproduction, including humans, where envelope protein genes of integrated endogenous retroviruses encode syncytins that mediate cell-to-cell fusion in placental tissues (Henzy and Johnson, 2013; Mi et al., 2000). Thus, elucidation of retroviral Env structures is relevant for the understanding of both virus-host interactions and basic physiologic mechanisms relevant to human health.

The Env of orthoretroviruses is cleaved by host cell proteases to form two independent subunits, the surface (SU) and transmembrane (TM) subunits, which remain covalently or non-covalently associated (Hogan and Johnson, 2023). The SU subunit binds host-cell receptors whereas TM mediates membrane fusion. The native Env on virions is composed of trimers of SU-TM heterodimers. The native trimeric Env is metastable, with TM assuming significantly different conformations in the pre-fusion and post-fusion states (Hogan and Johnson, 2023). The SU (gp120) and TM (gp41) subunits of the human immunodeficiency type 1 virus (HIV-1) pre-fusion Env trimer are closely associated, with the TM subunit assuming a relatively complex metastable helical fold (Pancera et al., 2014). In contrast, in the post-fusion state the SU and TM subunits are dissociated, with TM assuming a stable six-helix coiled-coil conformation (Chan et al., 1997). The transition between the two states is triggered by receptor binding. Following receptor binding, the fusion peptide (FP) in the amino-terminus of TM inserts itself into the target cell membrane and TM refolds itself into the six-helix bundle state, bringing the two membranes together to induce fusion.

Retroviral Env is classified into two major groups according to the manner by which SU and TM interact in the Env trimer (Henzy and Coffin, 2013; Henzy and Johnson, 2013; Hogan and Johnson, 2023). The major Env group is the gamma-type, of which the Env of gammaretroviruses is the prototype but also including the Env of alpharetroviruses, where SU and TM are covalently linked by an intersubunit disulfide bond (Hogan and Johnson, 2023; Leamnson and Halpern, 1976; Pike et al., 2011; Pinter et al., 1997). The third cysteine residue in a conserved CX_n_CC motif of TM forms this intersubunit disulfide bond whereas the first and second Cys residues form a disulfide loop within TM. The cysteine in SU forming the intersubunit disulfide bond has a more variable context depending on retroviral group. In the gammaretroviruses, the second Cys residue of a conserved CXXC motif forms the intersubunit disulfide bond whereas in the alpharetroviruses an amino-terminal Cys residue has that function (Pike et al., 2011; Pinter et al., 1997; Sjöberg et al., 2006). Importantly, the GP2 protein of filoviral envelope spike proteins (SP), including Ebola virus (EBOV), share sequence similarity with the TM of retroviruses. In fact, the GP2 subunit of filoviral SP also has a CX_n_CC motif that forms a disulfide bond with an amino-terminal cysteine in the SU-equivalent GP1 as in gamma-type Env (Bénit et al., 2001; Gallaher, 1996; Jeffers et al., 2002). The second major retroviral Env class is the beta-type Env of betaretroviruses and lentiviruses (Henzy and Coffin, 2013; Henzy and Johnson, 2013). The beta-type Env lacks a covalent intersubunit disulfide bond and has a CX_n_C motif in TM instead.

Beyond the common features of retroviral Env described above, the Env of retroviruses is highly diverse. The most intensively studied retroviral Env outside the primate lentiviruses is that of the gammaretroviruses. A recent review describes the structure and function of gammaretroviral Env in detail (Hogan and Johnson, 2023). The most important difference between the Env of most gammaretroviruses relative to other retroviral groups is the segregation of SU into two independently-folding domains, the amino-terminal receptor-binding domain and a more conserved carboxy-terminal C-domain thought to interact with TM. These two domains are linked by a relatively long proline-rich region (PRR) that modulates Env fusion (Barnett et al., 2003, 2001; Battini et al., 1992; Fass et al., 1997; Pinter et al., 1997; Valsesia-Wittmann et al., 1997). In contrast, the two domains of HIV-1 gp120 are closely associated structurally, with both domains participating in receptor and co-receptor binding (Kwong et al., 1998).

The high sequence diversity of retroviral Env, especially in its SU region, has precluded more detailed understanding of its evolution. Comparison of Env protein structures, which should evolve more slowly than the underlying protein sequences, should extend our understanding of the evolution of retroviral Env. Unfortunately, Env structural information is highly skewed, with many high-resolution monomeric and trimeric HIV-1 Env structures but more limited information for the structure of Env from other retroviral groups. The latter encompass crystal structures for the receptor-binding domain of type-C gammaretroviruses, including the murine (MLV) and feline (FLV) leukemia viruses (Barnett et al., 2003; Fass et al., 1997), lower resolution Env electron density maps of trimeric MLV Env (Förster et al., 2005; Löving et al., 2012; Riedel et al., 2017; Wu et al., 2008), an endogenous syncytin-2 SU structure in complex with its receptor (Martinez-Molledo et al., 2022) and post-fusion TM helical bundle structures of several different retroviruses. However, these structures have not led to deeper insights into the evolution retroviral Env. Several reasons make determination of retroviral Env structures experimentally challenging. These include the metastable state of pre-fusion Env and the complex multidomain SU organization probably associated with conformational flexibility. These challenges will probably continue to hinder detailed understanding of the function of Env proteins in different retroviral groups as well as understanding of the evolution of this complex and highly diverse retroviral protein.

A significant breakthrough in structural biology is the emergence of protein structure prediction tools based on machine learning and artificial intelligence methods. In particular, AlphaFold2 (AF2) has been very successful in generating structural models of proteins with high confidence, including for proteins without structural precedent in the Protein Data Bank (PBD) (Jumper et al., 2021; Tunyasuvunakool et al., 2021). The broad availability of AF2 has provided an opportunity to extend the understanding of retroviral Env structure and function as well as its evolution.

AF2 is a neural network-based tool for protein structure prediction (Jumper et al., 2021). The AF2 neural network was trained on protein structures and their sequences present in the PDB until about 2020. While AF2 can make use of templates for modeling, these are not necessary for structural modeling. This *ab initio* modeling can be successfully performed even when similar structures are not present among those used for AF2 training. Importantly, AF2 reports quality scores for its final and intermediate models, allowing interpretation of model reliability and selection of models among those in the output. Highlighting the ability of AF2 to generate unprecedented retroviral Env structural models *ab initio* with high reliability, a model of the simian foamy retrovirus (SFV) SU, which had no precedent in the Protein Data Bank (PDB) at the time it was generated, aligns closely to subsequently obtained experimental crystal structures of SFV SU, with the main deviations in the alignments corresponding to the low-scoring regions in the AF2 model (Fernández et al., 2023; Hötzel, 2022).

AF2 has been applied to the modeling of SU structures of lentiviruses and betaretroviruses (Hötzel, 2021). The SU structural models yielded both expected and unexpected results. Among the expected results was the confirmation of a partial structural similarity between the SU of the beta-type betaretroviruses and lentiviruses limited to a TM-proximal region of SU, named the proximal domain (PD), formed by a complex β-sandwich structure, in agreement with previously determined fragmentary sequence similarities between the SU of these two retroviral groups (Hötzel, 2021, 2008; Hötzel and Cheevers, 2001). The structural models further indicated that the apparent structural diversity of beta-type lentiviral and betaretroviral SU, including its domain configuration, is the result of differential expansion of loops in the conserved PD in the betaretroviruses and lentiviruses (Hötzel, 2021).

Subsequent application of AF2 to the modeling of the SU of other retroviral groups yielded additional unexpected results. Modeling of the SU of gammaretroviruses revealed that most of the C-domain is comprised of a PD with the same overall β-sandwich structure as the PD of the betaretroviruses and lentiviruses without significant internal expansions (Hötzel, 2022). The RBD and PRR of gammaretroviral SU is an amino-terminal extension of the conserved PD. In addition, the Env of some gammaretroviral and alpharetroviral endogenous elements are comprised of a lone PD without an amino-terminal RBD, including some without any significant expansions and others with relatively long internal PD expansions. Structural modeling of retroviral SU has provided significant insights into the evolution of this protein family, identifying conserved structural domains that are not obvious from sequence alignments and mechanisms of structural diversification based on expansion of the conserved PD.

Trimeric Env can potentially provide additional insights into the evolution of retroviral Env. After the PD was identified as a universal retroviral SU structure it became evident that the gamma-type SP of filoviruses also has a PD structure in GP1 in the same general location as the PD of retroviruses, closely associated to GP2 in SP trimers (Hötzel, 2022). Remarkably, the PD of filoviruses is oriented significantly different in SP trimers compared to the PD of HIV-1 in Env trimers, with a shift of about 120° around the vertical trimer axis. This results in different GP1 PD regions interacting with GP2 compared to the contacts the PD of HIV-1 gp120 makes with gp41. In addition, the organization of GP2 and gp41 in pre-fusion trimers also differ significantly structurally (Lee et al., 2008; Pancera et al., 2014). Analysis of glycosylation site patterns of gammaretroviral SU suggest that the orientation of the PD in the trimeric Env of gammaretroviruses resembles that of filoviruses rather than the lentiviruses (Hötzel, 2022).

A more recent variant of AF2, AF2 multimer, allows modeling of multimeric protein complexes (Evans et al., 2021), providing an opportunity compare the trimeric Env of different retroviral groups structurally. Here, AF2 multimer was used to model the trimeric Env of the prototype type-C gammaretrovirus, MLV, the Env of an endogenous gammaretroviral element in the RDR interference group (RDR Env), and the variant gamma-type Env of avian leukemia alpharetrovirus (ALV). These trimeric Env models were compared with existing trimeric HIV-1 Env and EBOV SP crystal structures to understand the evolution of trimeric retroviral Env. The trimeric Env models confirm the close structural relationship between gamma-type Env and filoviral SP, with the development of novel intersubunit interactions in the gammaretroviruses, providing insights into the evolution of trimeric gammaretroviral Env.

## RESULTS

### Modeling of trimeric MLV Env with AF2 multimer

The AlphaFold Structure Database (AFDB) (Varadi et al., 2021) includes structural models for a large number of endogenous, uncleaved Env elements from multiple vertebrate species of varying quality. However, multimeric protein models, including trimeric retroviral Env, are not yet part of the AFDB. Therefore, structural modeling of trimeric Env was performed here using AF2 multimer.

Selection and vetting of models were guided by the quality and reliability scores yielded by AF2 and by visual examination of models to exclude those with artifacts incompatible with previously described biochemical data. AF2 and AF2 multimer has four metrics for structural model reliability (Evans et al., 2021; Tunyasuvunakool et al., 2021). One is a metric of local structural confidence, the predicted local distance difference test (pLDDT), which ranges from 0 to 100. In practice, pLDDT scores of 90 or above indicate high side-chain conformation confidence, whereas scores of 70 and above indicate good main-chain reliability. Scores between 50 and 70 reflect lower confidence and, below 50, poor confidence or effectively unfolded states. The second metric is the predicted template modelling (pTM) score, a more global metric of model reliability, with pTM scores above 0.5 (in a scale from 0 to 1) indicating similar fold. A related metric is the interprotomer pTM, or ipTM, which provides a similar global metric for reliability between different protomers in the model. The fourth metric is the positional alignment error (PAE). This is calculated on a per-residue manner, yielding a matrix where the predicted error, in Å, in the position of residue *x* in models aligned on residue *y*. This metric is useful to identify domains and subdomains with variable relative position or interdomain flexibility.

Several modeling strategies were explored in preliminary trimeric MLV Env modeling runs, including control modeling runs to interpret the role of different input parameters and potential biases embedded in the AF2 neural networks. These preliminary modeling runs were performed with the Env sequences of different MLV strains with different linker sequence lengths between the SU and TM regions, including different subsets of the membrane-spanning domain (MSD) and intraviral (IV) TM regions and different templating strategies. Structural models with low quality scores, TM in conformations resembling post-fusion states, unfolded or with different types of domain swaps in which the polypeptide chains in the TM regions cross the trimer axis into an adjacent Env protomer (defined here as a single SU/TM heterodimer) and incompatible with MLV Env intersubunit disulfide biochemical data (Löving et al., 2012) were not further pursued. A stepwise modeling strategy successfully yielding a mature MLV Env trimer structural model is described below and summarized in Figure 1A.

**Figure 1.**
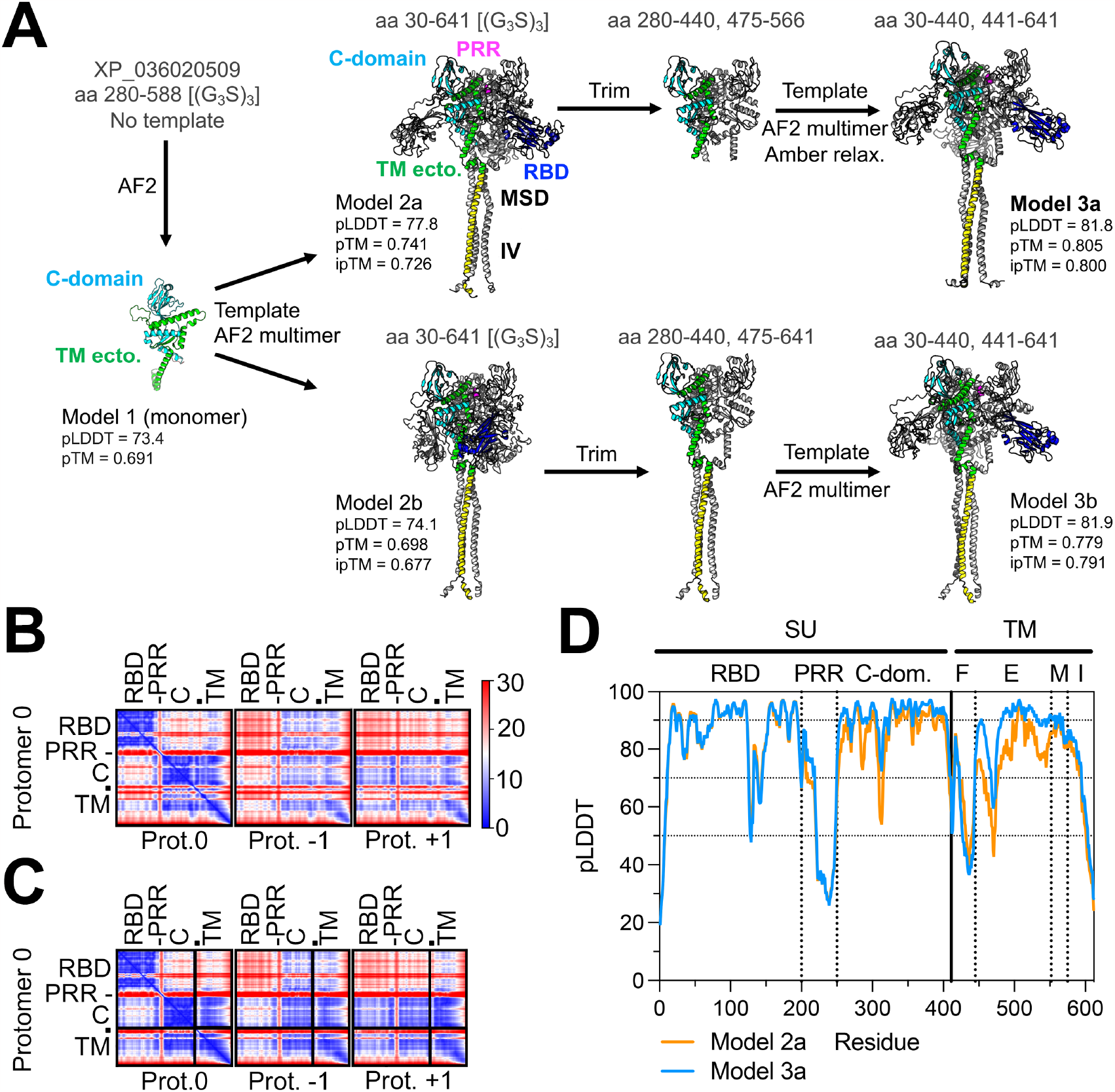
Trimeric Env modeling strategy and quality scores. (A) Modeling of trimeric MLV Env, showing intermediate models and templates used for each step. The global pTM and ipTM and average local pLDDT quality scores are shown for each model. The amino acid sequence boundaries for each model and template are indicated above the model vignettes. The vignettes visually summarize the structural boundaries of the models and templates. The approximate location of the SU and TM regions are indicated in the model 2a vignette. The [(G_3_S)_3_] notation indicates the linker added between residues 411 and 412 in models 1, 2a and 2b. Amber relaxation was only performed for model 3a. The “TM ecto”, “MSD” and “IV” labels indicate the ectodomain, membrane spanning domain and intraviral regions of TM. (B) PAE scores trimeric Env model 2a. The leftmost panel indicates PAE scores within each protomer (“Protomer 0”), with the approximate location of the main Env regions shown on the left and top of the matrix. The dot indicates the location of the fusion peptide in the matrix and C indicates the C-domain of SU. The middle and right panels show the PAE of “protomer 0” relative to the protomers in the counterclockwise (−1) and clockwise (+1) directions as observed from the top of the trimer. The PAE scale, in Å, is shown to the right of the panels. The other effectively redundant matrices for each of the two other protomers in the trimer are not shown for simplicity. (C) PAE matrix for model 3a, as shown in (B). (D) Positional pLDDT scores of models 2a and 3a. The boundaries or the major SU and TM regions are shown by vertical dotted lines. F, E, M, I indicate the fusion peptide, ectodomain, membrane-spanning domain and intraviral regions of TM. The threshold pLDDT scores of 50, 70 and 90 are shown by dotted lines.

An AF2 structural model of the C-domain of SU and TM ectodomain from a polytropic MLV strain joined by a (G_3_S)_3_ linker sequence at the Env cleavage site was obtained without templates. The resulting model was very similar to a polytropic Env model in the AFDB (UniProt accession P10404) but with slightly better pLDDT scores in the TM region. This monomeric Env fragment model was then used as a template for full-length trimeric Env modeling including a (G_3_S)_3_ linker between SU and TM in two independent runs at different dates to account for changes in random seeds and the stochastic nature of AF2 structural modeling. The top-scoring trimeric Env models 2a and 2b from each run were selected for further analysis and modeling. The main difference between the monomeric Env fragment template and the trimeric Env models was observed at the top and center of the trimer, where a kinked helix corresponding to the center of the trimer in the monomeric Env template is observed as a longer straight α-helix in the trimeric models. In addition, although the PD and TM regions and the relative position of different elements are similar in different models, RBD position varies (compare overall shape of models 2a and 2b in Figure 1A vignettes for an overall view). These correlations between PD and TM regions within and between protomers and the lack of correlation between the RBDs and other parts or the trimer are reflected in the PAE scoring matrices (Fig. 1B). Local pLDDT scores of the top structural models were relatively high except in the PRR, fusion peptide, a short region at the top and center of the trimer in TM and in the intraviral region of TM (Fig. 1D). Visual inspection of the models revealed that the (G_3_S)_3_ linker between SU and TM is sandwiched between the RBDs and TM regions in slightly different conformations in different models, potentially interfering with RBD positioning. Thus, a final round of modeling of cleaved, mature trimeric Env without linkers was performed.

Modeling of mature trimeric Env was performed by providing the SU and TM sequences separately. Modeling using the monomeric Env Model 1 as template resulted in TM domain-swapped models only. Therefore, sections of the trimeric models 2a and 2b were used as templates. The first model used only the C-domain and TM ectodomain regions of the trimer in model 2a as template, without the signal sequence, (G_3_S)_3_ linker and TM membrane-proximal (MPER) regions as template. The template for an independent modeling run had the same regions of model 2a-based template but also included the MSD and IV sections of TM. The top-scoring models 3a and 3b from this second round from each different template converged on highly similar structures within the PD and TM regions, differing in those regions only by minor shifts in the positions of some helices in the TM region ectodomain. However, the positioning of the RBDs relative to the rest of the trimer remained variable even without the linker sequence, although in a generally similar orientation among models, indicating that the variations in RBD position in the models were not due to artifacts introduced by linker regions and represent true uncertainty in the models. The RBD position uncertainty and the higher correlation between the PD and TM regions of the trimers are reflected in the PAE scoring matrices for the final model 3a (Fig. 1C). The reliability scores of models 3a and 3b were generally higher than for the first round (Fig. 1A-D), with the caveat that these may be relatively overconfident due to high template and model sequence identity (Roney and Ovchinnikov, 2022). However, the convergence of top-scoring models 3a and 3b obtained from slightly different templates indicates the mature trimeric Env models as probably being better than the preceding models with the (G_3_S)_3_ linker.

The top-scoring model 3a from the third round was further processed by amber relaxation to resolve residue clashes. The major differences between the relaxed and unrelaxed models were in residues Glu-481, Leu-464 and Thr-465 (numbering from the first residue of the Env model, residue 30 in XP_036020509) in the turn between two TM α-helices close to the distal end of TM adjacent to the trimer axis and the nearby SU His-314. Extensive clashes are observed among these residues from different protomers in the unrelaxed model whereas in the relaxed model many of these clashes are resolved by changes in side-chain rotameric conformation of these residues that break the local trimer symmetry in that region of the model. These TM residues are located in the region of the chain that crosses the trimer axis to form the major domain swap observed in some models. This suggests that the artifactual domain swaps in several trimeric Env models are a consequence of local clashes among axis-proximal residues that arise from symmetrical modeling of the protomers by AF2 multimer. Analysis of relaxed model 3a using the MolProbity server (Davis et al., 2007) indicated good overall model structure quality, with 36 unique residue clashes within and between chains (77 total in the three protomers) and 5 unique Ramachandran main-chain angle outliers (11 total). Importantly, clashes between protomers in the model are located mostly in the periphery of the trimer involving regions that are likely flexible, the PRR and fusion peptide, none in the hydrophobic inter-protomer interface in the trimer axis (see below).

It is possible that AF2 uses previously known viral and HIV-1 Env trimer structures in its training dataset as implicit templates for MLV Env trimer modeling that bias the resulting models, rather than yielding models closer to the true trimeric MLV Env structure. The most obvious implicit templates are EBOV SP structures. This was addressed by control modeling of the Zaire strain EBOV trimeric SP without templates, as it was done for MLV Env trimer modeling. Initial modeling of the GP1 base and head subdomains that comprise the PD of GP1 (Hötzel, 2022; Lee et al., 2008), without templates, resulted in unfolded models even though several GP1 structures were presumably included in AF2 training. Similar results were obtained when attempting to model monomeric GP1/GP2, even when using a GP1 crystal structure as a partial template. Models of the EBOV SP trimer in the pre-fusion state were only obtained when enabling automatic selection of templates from the PDB. These results indicate that it is unlikely that structural information embedded in AF2 was “recalled” to model the MLV Env trimer and the intermediate models used as templates. Thus, the trimeric MLV Env *ab initio* model obtained here arises from the sequence information within the input, the sequence diversity of retroviral Env sequences in the multiple sequence alignments used by AF2 to determine residue covariations and co-evolution, and protein folding information embedded in AF2.

Structural alignments of the MLV Env trimer model to PDB structures and models in the AFDB were used to assess its uniqueness. A single protomer of the trimeric polytropic MLV Env model excluding the RBDs and PRR was used in structural alignment searches to identify homologs in the PDB and the AFDB using the Foldseek server (Kempen et al., 2023). Although several monomeric endogenous gammaretroviral Env models in the AFDB were identified as close homologs, no structural homologs were identified in the PDB. The DALI Protein Structure Comparison server (Holm, 2020) identified, among 635 spurious hits unrelated to viral proteins, 3 viral envelope protein structures. In decreasing hit score order these were an EBOV SP trimer (PDB 6VKM), a post-fusion structure for a baculovirus gp64 envelope protein (PDB 3DUZ) and a trimeric human immunodeficiency virus type 1 (HIV-1) Env structure (PDB 6PWU). The latter aligned with the MLV Env trimer model mostly in the PD regions previously shown to be conserved among retroviruses, with very limited alignment in the TM regions. Therefore, the trimeric MLV Env structure is unprecedented in its details and distinct from previous known structures.

### SU structure in trimeric Env

The SU region in the MLV Env trimer is very similar to previously described structures and models. The RBD assumes the same L-shaped Ig-like fold as a previously described ecotropic MLV RBD crystal structure (PDB 1AOL) (Fass et al., 1997). The polytropic Env RBD model and structure aligned with a root-mean-squared deviation (rmsd) of 2.11 Å and TM-score of 0.70. As mentioned above, the RBD was located not at the top of the trimer in the final model or the best intermediate models, as might be expected, but rather on the sides of the trimer pointing towards the viral envelope membrane with varying angles in different models. Each RBD was associated with the PD and TM regions of the Env protomer in the counter-clockwise direction when observed from the top of the trimer, consistent with the cross-protomer PAE scores between the RBD and PD and TM regions (Fig. 1B and 1C, protomer -1 matrix). The orientation of the RBD along its main axis also varied among models. In most models the RBD variable regions A and B (VRA and VRB) (Barnett et al., 2003) point in the counter-clockwise direction when the trimer is observed from the top, but in some models the RBD is rotated about 180° along its main axis, resulting in the VRA and VRB regions pointing in a clockwise direction. In some lower-scoring models the RBDs are positioned on top of the trimer pointing away from the envelope membrane. The uncertainty about the RBD position is not unexpected given that the RBD is linked to the rest of the SU region by the relatively long PRR. Residues 203, 205, 208, 210, 211 and 214 in a more conserved region of the PRR are anchored by hydrophobic and hydrogen bond interactions in pockets within a larger crevice formed by TM and the PD near the center of the trimer axis (Fig. 2B and C). The rest of the PRR from that anchor region to the C-domain assumes an unfolded conformation with low pLDDT scores in all models (Fig. 1D, 2A and 2B).

**Figure 2.**
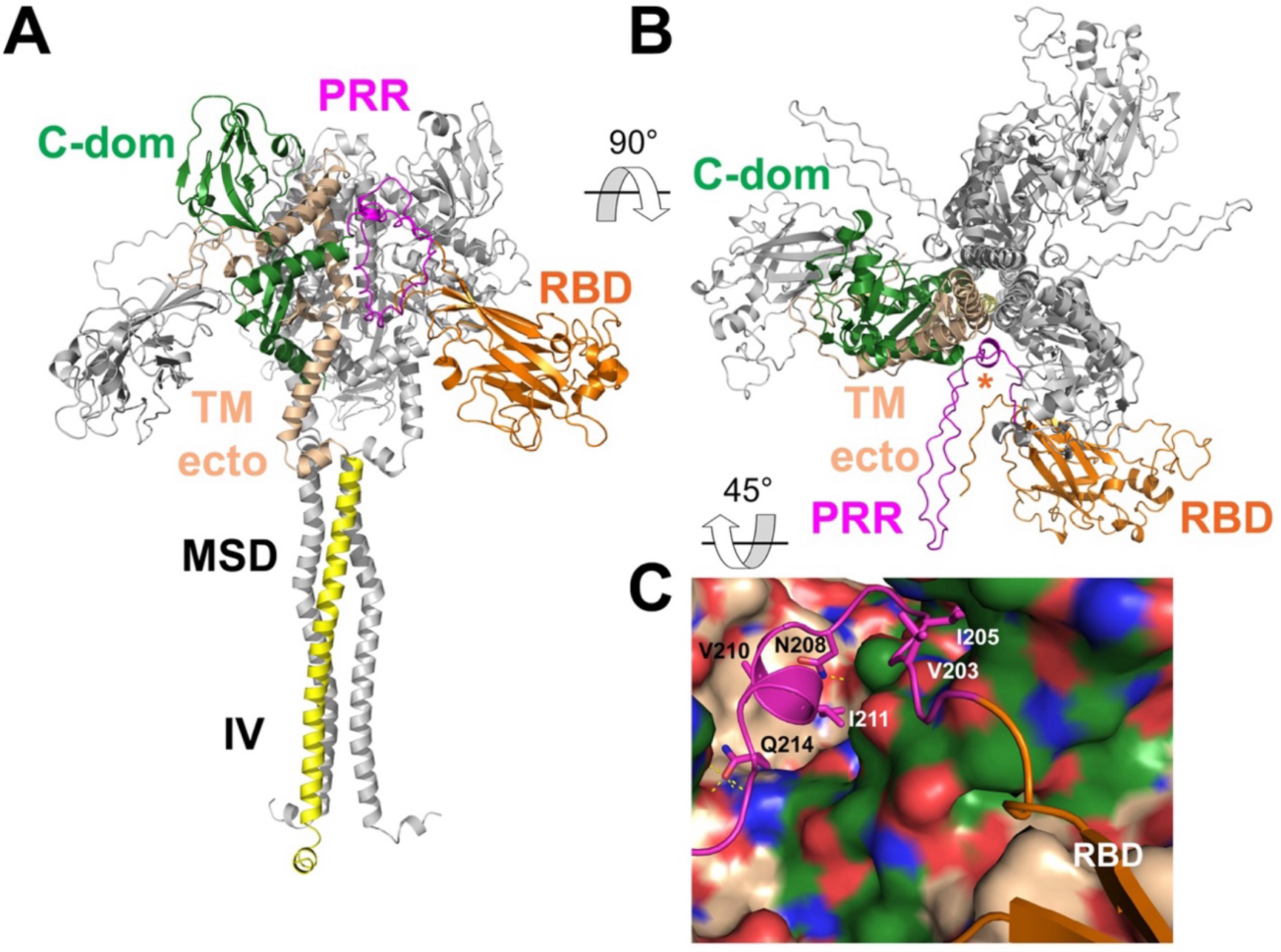
Overall view of the trimeric MLV Env model and PRR anchoring. (A) Lateral view of trimeric MLV Env model 3a. The regions of a single protomer are shown in color for clarity, with color-coded labels. MSD and IV indicate the membrane-spanning domain and intraviral regions of TM, in yellow. (B) Top view of the trimeric MLV Env model, in the same color scheme as in (A). The orange asterisk indicates the helical PRR anchor along the central TM helices. (C) detailed view of the anchoring residues of the PRR (magenta), with the TM (salmon) and C-domain (green) regions shown in surface rendering. Red and blue areas indicate negative and positive charges. The RBD preceding the PRR is shown in orange. Dotted lines indicate potential hydrogen bonds. The rotation angles and directions from panels (A) to (B) and (B) to (C) (approximate) are indicated.

Unsurprisingly, the PD region is also essentially identical to the previously reported monomeric MLV PD AF2 model (Hötzel, 2022), without any significant differences due to the trimeric context. As described below, the C-domain section between the PRR and PD regions forms an α-helix (αPD1) that is closely associated with TM and the terminal PD sections rather than the core PD regions. In addition, the C-terminus of SU also forms an α-helix (αPD5) closely associated with TM (Fig. 2A and 3C).

### Pre-fusion MLV TM assumes a conformation similar to pre-fusion filoviral GP2 in trimers

The TM ectodomain regions form the center of the trimer model along its vertical axis in a mostly helical structure. The MLV Env trimer has four α-helices in the TM ectodomain, designated here αTM1 to αTM4 (Fig. 3C and G). The αTM2 helices from different protomers are positioned in a parallel configuration around the vertical trimer axis and form extensive, mostly hydrophobic, interprotomer contacts along the axis. The central αTM2 helices align with an rmsd of 0.407 Å to the corresponding regions of a xenotropic MLV TM structure in the post-fusion state (PDB 4JGS) (Aydin et al., 2013) even though that structure was not used as template, suggesting that inter-protomer interactions in this region remain essentially unchanged in the fusion process. Helix αTM2 includes all 17-aa residues of an immunosuppressive domain preceding the CX_6_CC motif (Bénit et al., 2001; Cianciolo et al., 1985; Henzy and Johnson, 2013; Hogan and Johnson, 2023), which remains fully occluded in the pre-fusion state. Its location in the central axis of the TM trimer in both the pre- and post-fusion states partially explains the high conservation of this region. Helix αTM2 is preceded by helix αTM1, which projects from the periphery towards the center of the trimer in an approximately 60° angle relative to the vertical axis. Helix αTM1 is in turn preceded by the fusion peptide. Helices αTM1 and αTM2 in pre-fusion TM form the first contiguous helix of the post-fusion TM coiled-coil. Helix αTM2 is followed by a loop including the CX_6_CC motif and helix αTM3. The αTM3 helices from different protomers are displaced away from the vertical trimer axis, resting on the αPD5 helices formed by the last residues of SU before the cleavage site (Fig. 3C). The last SU residue before the cleavage site is positioned close to the trimer axis, implying that Env cleavage must occur prior to final assembly of that region of the Env trimer.

**Figure 3.**
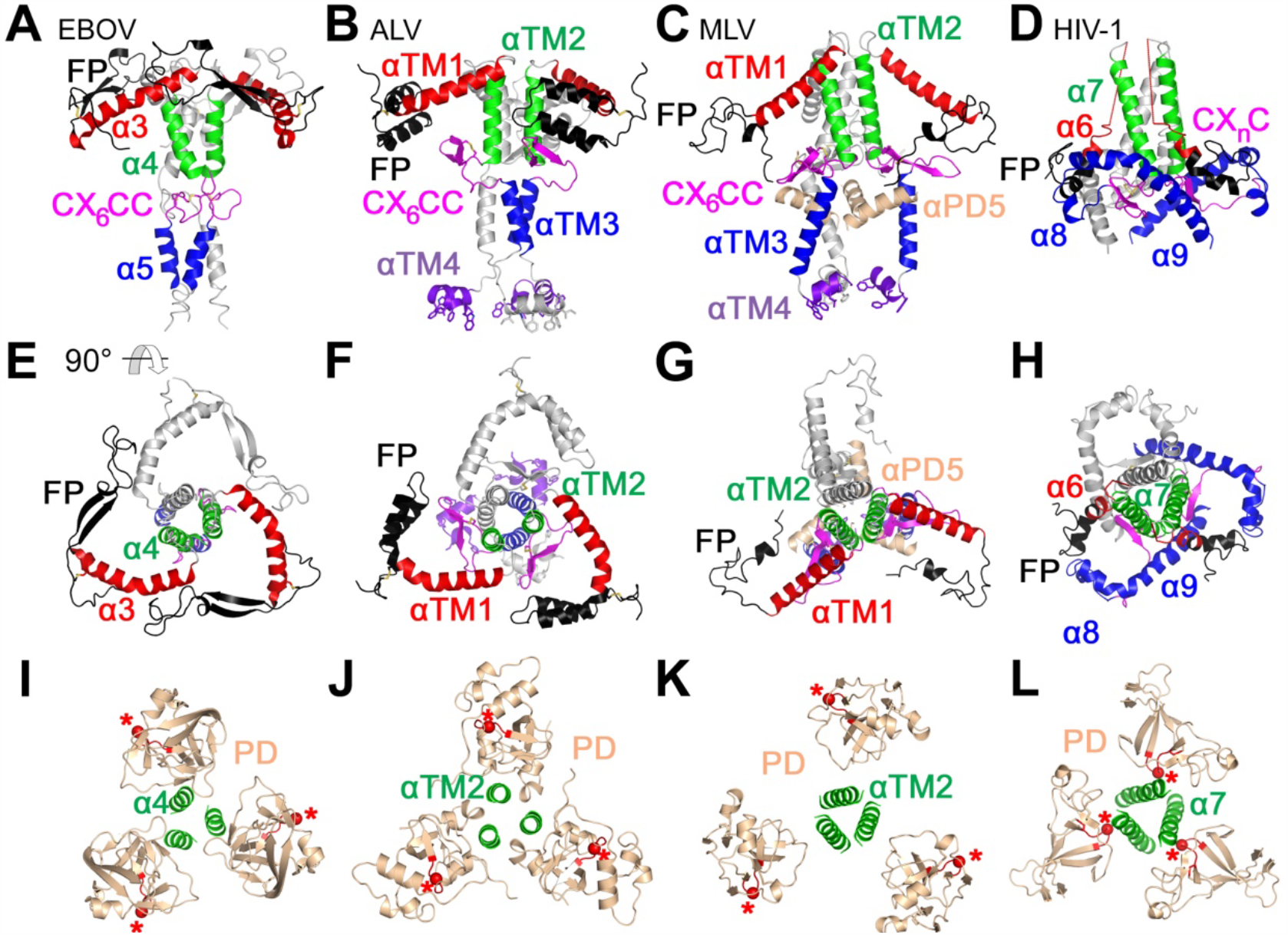
Structural similarity between pre-fusion gammaretroviral TM and filoviral GP2. The structures and structural models of the pre-fusion, trimeric EBOV GP2 and retroviral TM are shown in panels A-H, with the lateral (panels A-D) and top (panels E-H) views shown. The viral membrane is located towards the bottom of the figures in the lateral views. The MSD and IV regions are omitted. The virus for the structures and models in each column (e.g., panels A, E and I) are indicated in panels A-D. The homologous GP2 and TM helices and regions are shown in the same colors, with color-coded labels. Only two of the protomers are colored for clarity, with the third protomer shown in gray. Panels I to L show the orientation of SU and GP1 PD in the trimeric structures and models from a top view as in panels E-H. Only the αTM2 homologues are shown for the TM and GP2 regions. The red asterisks show the location of the PD turn between the conserved β-strands β4 and β5 (Hötzel, 2022). Note the outward-facing orientation of the β4/β5 turn for the EBOV SP and ALV and MLV Env models and the inward-facing orientation of the same turn in the HIV-1 trimeric Env structure. Note also the large hydrophobic side-chains of αTM4 (shown as sticks) in the MLV and ALV models that project towards the envelope membrane in panels B and C. EBOV SP structures, PDB 5JQ3. HIV-1 Env structures, PDB 4TVP.

The overall configuration of the αTM1, αTM2 and αTM3 helices is similar to that of corresponding α-helices α3, α4 and α5 in the GP2 of the EBOV SP trimer (Zhao et al., 2016), but different from the α-helices of HIV-1 gp41 except for the central α-helix α7 that corresponds to MLV Env αTM2 and EBOV SP α4 (Fig. 3A, E, D and H). As previously expected from the analysis of gammaretroviral SU glycosylation patterns, the orientation of the PD in the MLV Env trimer model is essentially the same as the PD of EBOV GP2 in the trimeric structures and different from the orientation of the HIV-1 PD in Env trimers (Fig. 3I, K and L), confirming the closer structural similarity between trimeric MLV Env and EBOV SP trimers that includes both organization of helical regions in TM and SU orientation.

The short amphipathic α-helix αTM4 forms an MPER that lies nearly parallel to the presumed envelope membrane, projecting large hydrophobic side-chains into the membrane plane as judged from the position of the MSD in the model just after αTM4 (Fig. 2A and 3C). Thus, the MPER extends the hydrophobic surface of the MSD. A similar helix-turn-helix structure has been described for the EBOV SP MPER/MSD region (Lee et al., 2017). An amphipathic α-helix in the influenza virus M2 protein partially embedded in the intracellular side of the membrane has been shown to enrich the M2 protein to the neck of budding virions by sensing membrane curvature and to promote the membrane lipid phase transitions necessary for virion release (Martyna et al., 2016; Schmidt et al., 2013). The topologically opposite MLV MPER may thus promote enrichment of Env on budding virions and facilitate the last steps of membrane fusion during infection. The N-linked glycosylation site of TM is located in the MPER αTM4 close to the trimer axis, with the side-chain of the asparagine pointing in the distal direction, away from the membrane. The MSD and IV regions of TM form contiguous α-helices that are closely associated among the protomers along the trimer axis (Fig. 2A).

### A unique structure in the MLV Env trimer anchoring SU to TM

Despite the overall similarity between the trimeric MLV TM model and EBOV GP2 structures, there are significant structural differences between the structural elements that anchor MLV SU to TM and EBOV GP1 to GP2 in the trimeric pre-fusion state. EBOV GP1 anchors in a hydrophobic structure formed by GP2, in a manner analogous to the anchoring of gp120 anchor to gp41 (Pancera et al., 2014). The SU anchoring structure in the MLV Env trimer model is similar to the one observed in the EBOV SP trimer but with additional structural elements unique to MLV Env that extend the anchoring structure in a manner similar, although not homologous, to HIV-1 Env. The main anchoring elements for the EBOV GP1 PD β-strands 2 and 11 homologues are GP2 α-helices α3 and α4 and parts of the fusion peptide (Fig. 4A and E). Similar interactions are observed in the MLV Env trimer model, with PD β-strands 2 and 11 contacting the GP2 α3 homologue, αTM1, and the fusion peptide (Fig. 4C and G). However, unlike EBOV GP1 and similar to HIV-1 gp120, the MLV PD region of SU has the β-strands 1 and 12 extensions emanating from the core region of the PD in a virion-proximal direction (Fig. 4C and G). These two β-strands, as the homologous β-strands in the trimeric HIV-1 Env structure (Fig 4D and H), are almost entirely embedded in a hydrophobic shell in the MLV Env trimer model. Several elements form this hydrophobic shell, including the fusion peptide, αTM2 and αTM3 in TM and the SU C-domain helix αPD1, which is positioned on the virion-proximal side of αTM1, and αPD5. An amino-terminal GP1 β-hairpin occupies a similar space in the EBOV SP trimers as αPD1 in trimeric MLV Env. The HIV-1 Env trimer does not have a structure homologous to MLV αPD1. The region occupied by αPD1 in MLV Env is instead occupied by gp41 helix α9 in HIV-1 Env (Fig. 4D and H). Thus, MLV Env has an SU anchoring structure that extends interactions homologous to those of EBOV SP in the proximal direction to include PD β-strands 1 and 12, similar to HIV-1 Env. However, the structures forming the virion-proximal section of the hydrophobic anchoring structure around PD β-strands 1 and 12 differ between the MLV Env trimer model and HIV-1 Env trimer structures, including a major MLV SU α-helix in that structure that is not observed in HIV-1 gp120.

**Figure 4.**
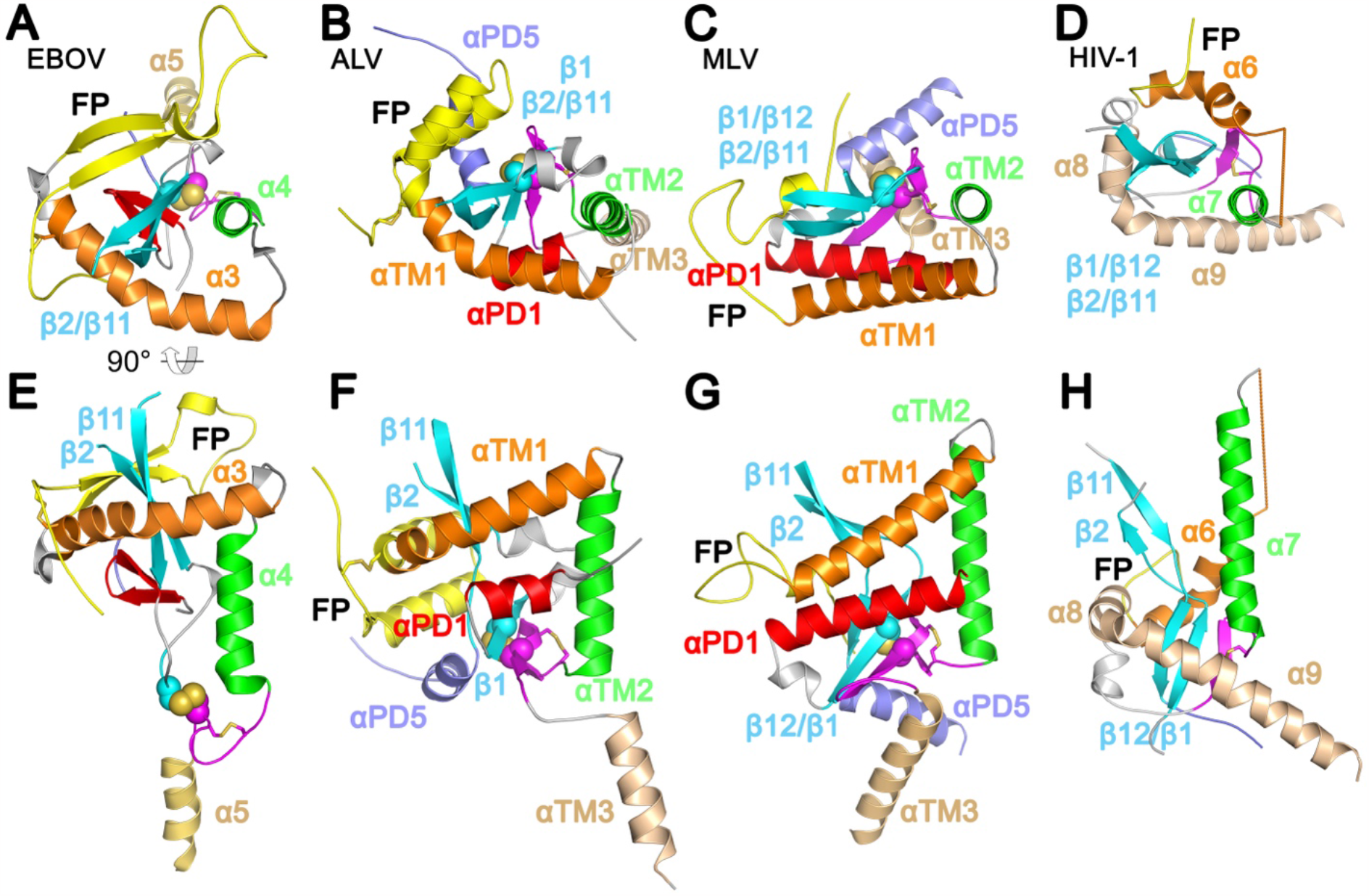
Structure of the hydrophobic SU and GP1 anchoring structures in retroviral Env and filoviral SP. Panels A-D show a top view of the anchoring structures for the different viruses as indicated in the panels. Only one protomer of the SP and Env structures and models is shown for clarity. The α4, αTM2 and α7 homologues in green are located in the center of the trimeric structures and models. Panels E-H shown the corresponding lateral view of the structures, rotated relative to the top panels as indicated in panel A, with the viral envelope towards the bottom of the figure. The homologous TM and GP2 structures are shown in the same colors in different panels and indicated in color-coded labels. The terminal PD β-strands of GP1 and SU are shown in cyan. The Cys residues mediating the intersubunit disulfide bonds in EBOV SP and ALV and MLV Env are shown as spheres. The straight line connecting α6 and α7 in the HIV-1 structure indicates an unresolved region of the structure. PDB structures as in Figure 3.

### Structure of the intersubunit covalent bond in the MLV Env trimer model

A major defining feature of gamma-type Env is the covalent disulfide bond between SU and TM and, in the gammaretroviruses specifically, the SU CXXC motif (CWLC in MLV) mediating this interaction. The MLV Env CWLC motif is located in PD β-strand 1, which is almost completely buried within the virion-proximal anchoring hydrophobic structure (Fig. 4G and 5A). The second Cys residue of this motif forms a disulfide bond with the third Cys residue of the CX_6_CC motif of TM within the same Env protomer in the trimeric model (Fig. 5A), consistent with previous biochemical data (Löving et al., 2012; Sjöberg et al., 2006). The first and second Cys residues of the CX_6_CC motif in TM form the expected disulfide loop near the trimer axis in the Env model (Fig. 5A). The two last Cys residues in the CX_6_CC motif are part of a β-strand that forms an extended β-sheet with PD β-strands 1 and 12 and a fourth β-strand within the TM cysteine loop (Fig. 5A). As expected, the first Cys residue of the CWLC motif, which is also entirely buried within the hydrophobic anchor, does not form a disulfide bond in the pre-fusion trimeric Env model (Fig. 5A). Thus, the isomerization of Env disulfides that occurs upon receptor binding required for cell infection (Hogan and Johnson, 2023; Wallin et al., 2005) must be preceded by significant conformational changes in SU and TM to expose both Cys residues in the CWLC motif. In contrast, the intersubunit disulfide bond between GP1 and GP2 in the EBOV SP is accessible on the surface of the trimer (Fig. 5C), thus defining another key structural difference between retroviral and filoviral gamma-type envelope proteins.

**Figure 5.**
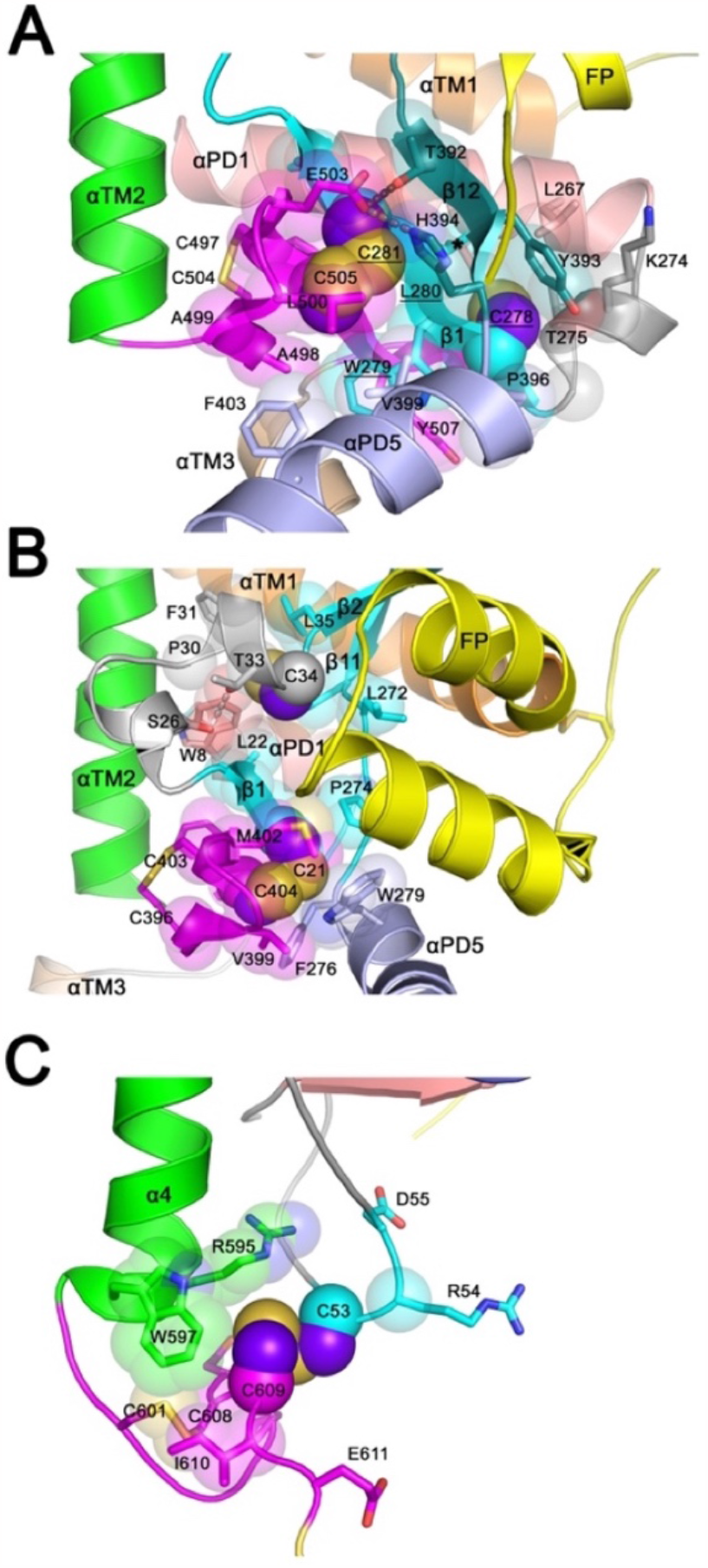
Structure of the intersubunit disulfide bond in trimeric gammaretroviral Env and EBOV SP. (A) MLV Env intersubunit disulfide. The disulfide bond between SU Cys-281 and TM Cys-505 is shown as spheres. Atoms surrounding the intersubunit disulfide are shown as semi-transparent spheres superposed on the corresponding residues shown as sticks and numbered. The different structural elements of TM and SU are shown in different colors and indicated with labels. The side-chain of unpaired Cys-278 in the CWLC motif is also shown as spheres with surrounding atoms shown as semi-transparent spheres. The model is shown from the side of one protomer with the viral envelope towards the bottom. (B) ALV Env intersubunit disulfide structure. The ALV Env model is shown in the same orientation as in panel A, with similar rendering and labeling scheme. The conserved unpaired Cys-34 is also shown as spheres. (C) EBOV SP intersubunit disulfide bond (PDB 5JQ3).

### Fitting of models into trimeric MLV Env electron density maps

The MLV Env model was assessed by fitting to previously described electron density maps of mature MLV Env. Several electron density maps have been described (Förster et al., 2005; Löving et al., 2012; Riedel et al., 2017; Wu et al., 2008). The best published map has a resolution of 15 Å and shows the Env trimer as a tripod-shaped structure with a region of low electron density in the center (Riedel et al., 2017). Four contiguous regions of high electron density in the periphery and top of the trimer were previously named the foot, heel, knee and head regions, starting at the base of the trimer (Fig. 6D). Previous attempts to fit the RBD structure to the Env electron density maps have put the RBD at the top of the trimer in a location expected to be available for receptor binding (Förster et al., 2005; Löving et al., 2012; Riedel et al., 2017).

**Figure 6.**
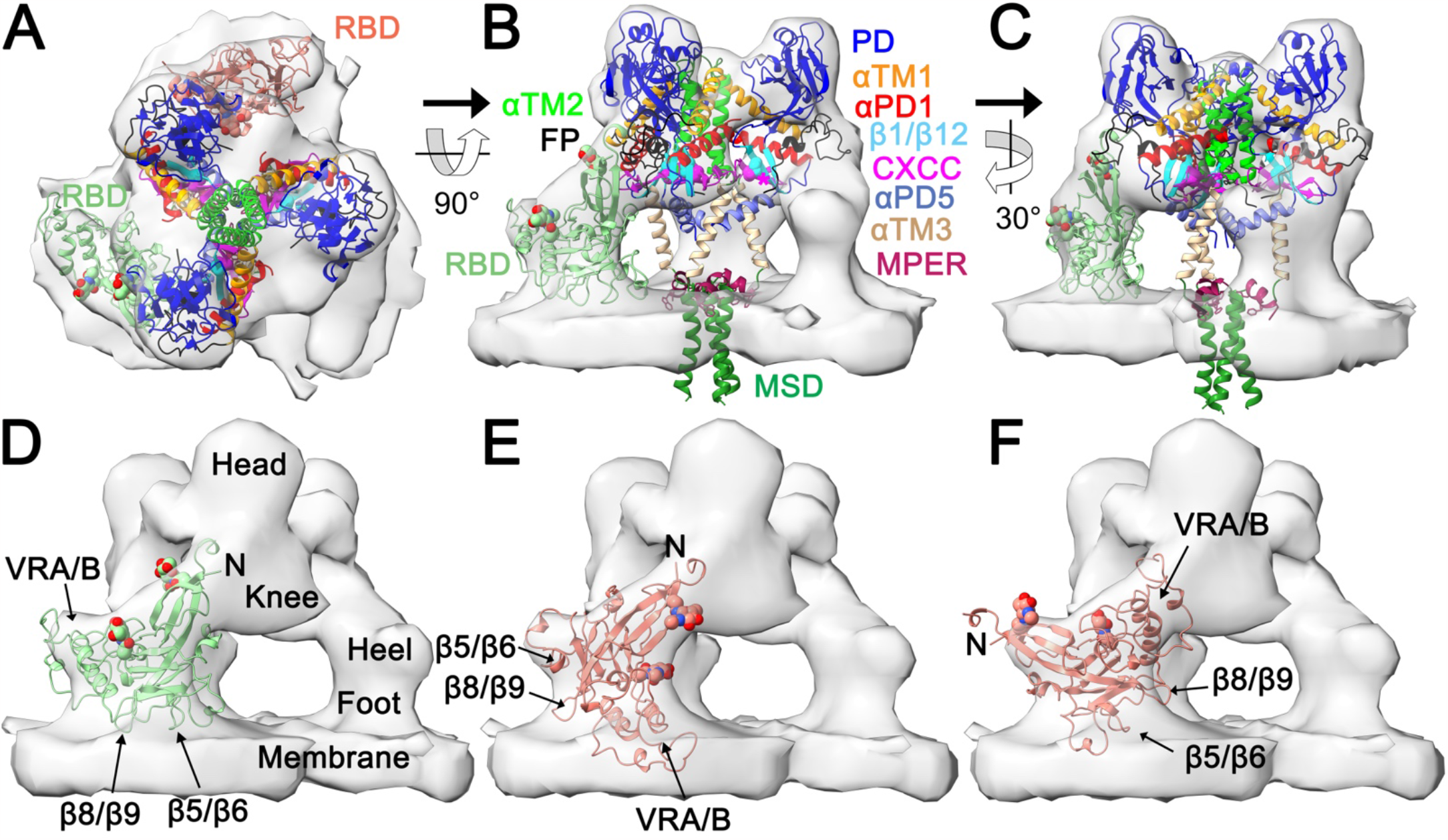
Fitting of the trimeric Env MLV model and RBD structure in the trimeric MLV Env electron-density map. Panels A to C show top (A) and side (B and C) views of the MLV Env trimer model without the RBD and PRR regions fitted to the composite trimeric MLV Env electron-density map (EMDB accession EMD-3373). Panel A also shows the structure of the MLV SU (PDB 1AOL) fitted in two different poses. The SU and TM sub-regions are shown in different colors indicated with color-coded labels. Panel (B) omits the second RBD pose shown in salmon in panel A for clarity. The structures and models are rotated as indicated between panels. The electron-density maps in panels D-F are rotated 60° to the right on the vertical axis relative to panel C to show alternative fittings of the RBD in the maps. The position of the VRA/B regions and loops between RBD β-strands 5/6 and 8/9 and the amino-terminus of SU (N) are indicated to show the relative RBD orientation. The location of the previously described head, knee, heel and foot regions of the electron-density maps are shown in panel D. The magenta asterisks in the middle section, right of center of panel B indicate the location of the intersubunit disulfide bonds shown as spheres in one protomer.

The MLV Env trimer model without the RBD and most of the PRR, which have low-confidence positions in the model, was fitted to the 15 Å resolution electron density MLV Env trimer map (EMDB accession EMD_3373) (Riedel et al., 2017) using ChimeraX (Goddard et al., 2017). The automatically optimized fit positioned the αTM4 helices and membrane-spanning regions of the model just above and within the membrane, respectively, and the SU PD region from strands β2 to β11 in the head regions of the maps at the top of the trimer (Fig. 6A-C). The central depression at the top of the trimer is lined by αTM1 and the distal end of αTM2 (Fig. 6A-C). The knee region includes the TM structure anchoring PD β-strands 1 and 12, with the CWLC motif and the inter-subunit disulfide bond in its core. The TM regions closer to the trimer axis are largely in the central regions of low electron density in the maps. However, a weak but clear electron density was described in the central region of some the MLV Env maps (Riedel et al., 2017). These lower-density map regions fit well visually with the central α-helices of TM and αPD5, including an intra-membrane density along the trimer axis that would correspond to the location of the membrane-spanning helices in the trimer model. The lack of strong density corresponding to the central TM regions at the resolutions attained for the MLV Env electron density maps was previously discussed (Riedel et al., 2017).

The only region in the electron density maps left for the RBDs based on these fitting results are in the heel and foot regions close to the viral membrane, away from the target cell. That is, the RBDs in the electron density maps appear to be in the same general position as the RBDs in the best quality AF2 Env trimer models. Indeed, that region of the electron density maps is L-shaped and should accommodate a similarly-shaped RBD. The MLV RBD crystal structure (PDB 1AOL) was manually fitted into that region while avoiding major obvious clashes with the PD and TM Env model regions fitted to the same maps (Fig. 6). The shape of the RBD and the heel and foot regions allow several possible fittings. In the first fitting the RBD was oriented with the VRA and VRB regions in the heel region pointing in the clockwise direction as seen from the top of the trimer (Fig. 6A-D, light green RBD moiety), in the opposite orientation as the RBD in MLV Env models 3a and 3b. In that orientation, the RBD touches the membrane and both RBD glycosylation sites point towards the outside of the trimer. In the second fitting, with the RBD turned 180° along its main axis relative to the first fitting, significant portions of the RBD embed in the membrane and one the RBD glycans points towards the center of the trimer (Fig. 6A and E, salmon RBD moiety). In a third alternative fitting, the RBD termini are located in the heel region, with the VRA/B regions pointing towards the knee region (Fig. 6F). In the last fitting, the RBD in the third fitting is turned around 180° along its main axis, resulting the VRA/B region being deeply embedded in the membrane (not shown) and, therefore, unplausible. Other possible arrangements, such as with the RBD termini in the foot region pointing towards the membrane (not shown), are also biologically and structurally unplausible. Although the first fitting with the VRA/B region located in the heel region is the most plausible and likely of the RBD poses, the shape of the RBD allows alternative fittings in the electron density map corresponding to the heel and foot regions that are not occupied by other Env model regions.

### Structural modeling of trimeric RDR Env

A second major gammaretroviral Env type is the RDR Env of the avian reticuloendotheliosis virus (REV), Mason-Pfizer monkey retrovirus (MPMV) and the related gammaretroviral endogenous elements RD114 and baboon endogenous virus (BaEV) (Hogan and Johnson, 2023; Sinha and Johnson, 2017). More distantly related endogenous members in the RDR receptor interference group include the human endogenous retrovirus W Env, or syncytin-1 (Syn-1) (Blond et al., 1999; Hogan and Johnson, 2023; Mi et al., 2000). Like MLV, the RDR Env has an amino-terminal RBD and a CXXC motif in SU. However, the SU of RDR group retroviruses and that of MLV and related gammaretroviruses have no significant sequence similarity besides the CXXC motif. In addition, the RBD of RDR Env models are structurally distinct from that of MLV and related gammaretroviruses, and are comprised of either a single or two tandemly repeated 3-strand β-sheets rather than the Ig domain-like fold of the MLV RBD (Hötzel, 2022). Trimeric RDR Env was modeled using the same overall strategy used for trimeric MLV Env to test whether the strategy for trimeric Env modeling can be generalized to other retroviruses and to understand how trimeric Env differs structurally between these two major gammaretroviral groups.

Several RDR Env sequences from exogenous viruses and endogenous elements were tested in preliminary modeling runs. High quality trimeric Env modeling was achieved with the Env sequence of an endogenous element from the black-and-white snub-nosed monkey *Rhinopithecus bieti* that is most closely related to the exogenous Env of simian retrovirus 2 and MPMV (74% and 68% amino acid identity over the entire mature Env sequence). Modeling of trimeric RDR Env required entering SU and TM sequences separately to allow resolution of the virion-proximal regions of the TM ectodomain. Thus, only two rounds of modeling were used, one for monomeric Env with a linker between subunits and a second round for trimeric cleaved Env. The AF2 quality scores of the final model were high (Fig. 7B, D and E) similar to the scores of second round trimeric MLV Env model 2a. The apical end of Env where helix αTM1 turns to helix αTM2 had relatively low pLDDT scores and obvious residue clashes in the unrelaxed models that were resolved by amber relaxation, similar to the trimeric MLV Env model. Despite the low pLDDT scores in that region, the turns between these helices are very similar in both models (Fig. 7F, compare to Fig. 4G). Analysis of the relaxed model using the MolProbity server indicated good overall model structure quality, with 37 unique residue clashes within and between chains (50 total in the three protomers) and 15 unique Ramachandran main-chain angle outliers (33 total).

**Figure 7.**
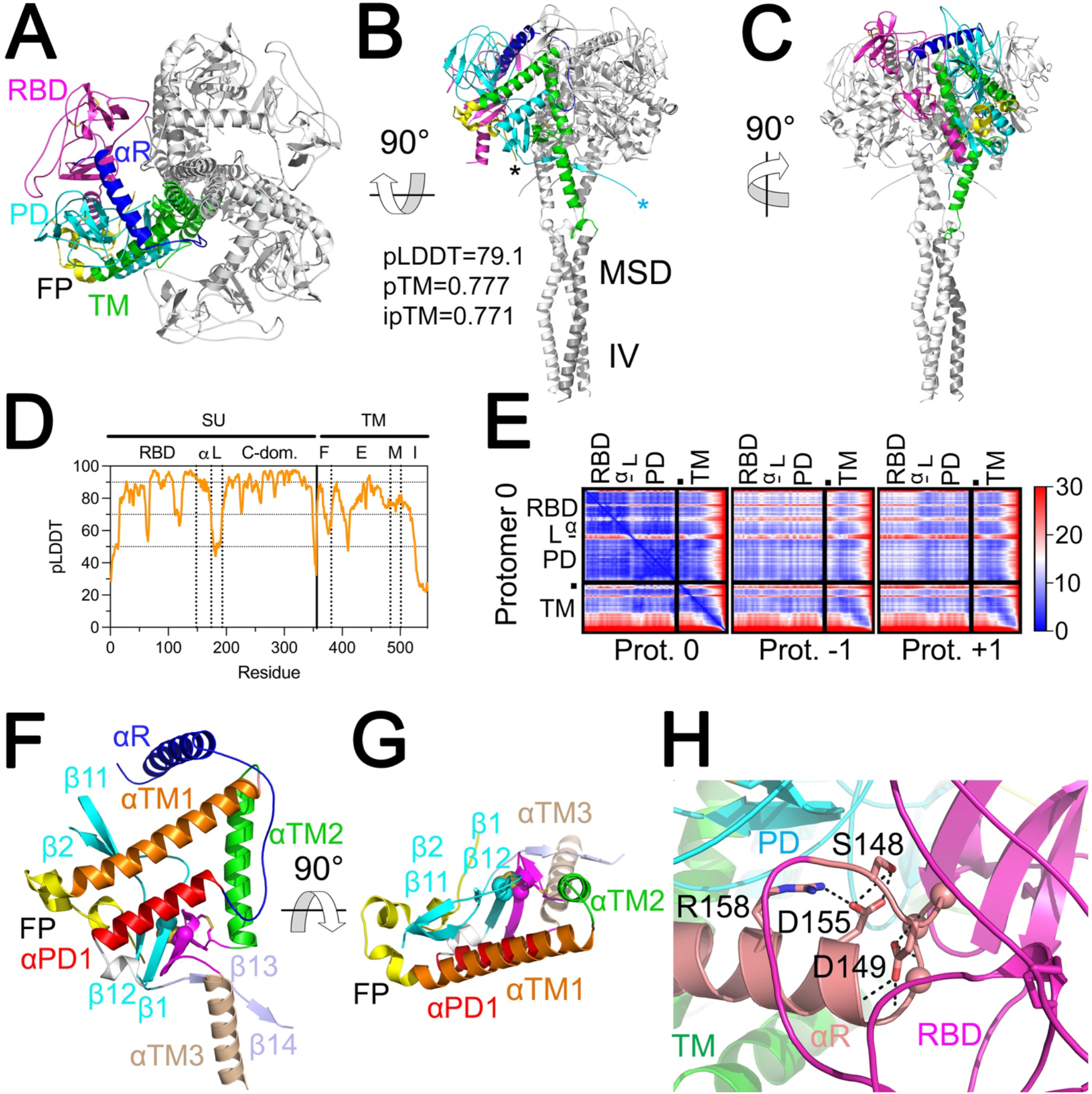
Structural model of trimeric RDR Env. Top (A) and lateral (B and C) view of the *R. bieti* trimeric endogenous RDR Env trimer model. The main sections of the Env ectodomain are indicated in different colors. Only one protomer is shown in colors for clarity. The models are rotated as indicated by arrows and angles between panels. The AF2 quality scores are shown in (B). The cyan and black asterisks indicate the C-terminus of SU and N-terminus of TM, respectively, within the same protomer. Note the long magenta RBD loop extending toward the PD contacting loop F on the left side of the model shown in panel (A). The last 22 disordered residues of TM in the intraviral region are not shown for clarity. (D) Positional pLDDT scores with Env sections indicated, separated by vertical lines. α, helix αR. L, RBD/PD linker region. (E) PAE scores of the trimeric RDR Env model. The leftmost panel indicates PAE scores within each protomer (“Protomer 0”). The middle and right panels show the PAE of “protomer 0” relative to the protomers in the counterclockwise (−1) and clockwise (+1) directions as observed from the top of the trimer. The approximate locations of the RBD, helix αR (α), linker region (L) and PD of SU and the fusion peptide and TM subunit are indicated on the axes. The PAE scale, in Å, is shown to the right of the panels. The other effectively redundant matrices for each of the two other protomers in the trimer are not shown for simplicity. Lateral (F) and top (G) views of the TM ectodomain region and associated SU regions showing the intersubunit disulfide bond as spheres. The αR is shown in panel F to highlight its position relative to the TM helical regions. The orientation of the model sections in panels F and G show the main SU and TM elements in similar orientations as Fig. 4G and 4C, respectively, following the same coloring scheme in these figures. (H) Detail of the structural model in the region encompassing the SDGGGX_2_DX_2_R motif between residues 148 and 158 of SU preceding and within helix αR (salmon). The conserved residue side-chains are shown as sticks except conserved Gly residue α carbons, shown as spheres. Polar interactions are shown as dashed lines.

Unlike the MLV Env model, the RDR RBD is more closely associated with the rest of SU in the same protomer by visual inspection of the model and assessment of the PAE score matrices (Fig. 7A and E). The RBD is positioned clockwise relative to the PD of the same subunit when the trimer is observed from the top (Fig. 7A), the opposite direction that is observed in the MLV model. The SU section of trimeric Env, including the RBD, aligns well with previous monomeric SU models of MPMV, BaEV, REV and RD114 (Hötzel, 2022), with rmsd ranging between 1.573 and 2.143 Å. Thus, high-reliability monomeric SU structural models are sufficient to capture the conformation of RDR SU in trimers, including RBD position within SU. In the trimeric Env model and the previous monomeric SU models a long loop between the second and third β-strand of the second 3-strand β-sheet of the RBD extends to contact the F loop in the apical region of the PD (Hötzel, 2022) in the same protomer (Fig. 7A). In addition, unlike the trimeric MLV Env model, the RDR RBD is exposed alongside the PD at the top of the trimer (Fig. 7A to C).

A significant difference between the trimeric MLV and RDR Env models is the structure and position of the linker between the main RBD section and the rest of SU. Unlike PRR of the MLV Env model, which extends laterally and is mostly unstructured, the linker between the RDR RBD and the PD is mediated by an α-helix (αR) located at the top of the trimer, followed by a shorter unstructured linker sequence connecting it to the αPD5 helix (Fig. 7A, B, C and E). This α-helix and the preceding loop region include one the most conserved regions of SU among RDR Env sequences, the SDGGGX_2_DX_2_R motif (Cheynet et al., 2006; Sinha and Johnson, 2017). This motif has been proposed to meditate receptor interactions. However, although this motif is located near the top of the trimer in the model, it is occluded by other RBD structures and would not be available for receptor binding without significant SU structural changes (Fig. 7H). Instead, the conserved residues in this motif mediate numerous polar interactions with each other and with the main chain to stabilize a turn between the main RBD structures and helix αR, with the three conserved glycine residues forming the core of the turn (Fig. 7G). The residues in the conserved motif (number 148 to 158 in the model), both within the loop and in αR, have high local pLDDT reliability scores of 90 or above (Fig. 7D), supporting the specifics of the intra-chain interactions in this region. The section of helix αR exposed at the top of the trimer, which may be required for receptor binding (Cheynet et al., 2006), starts at Glu-159 in the model, immediately after the conserved motif. The RBD at the periphery of the trimer between protomers and helix αR at the top of the trimer give the RDR Env trimer model a nearly flat shape at its distal end, unlike the turret-shaped distal end of the trimeric MLV Env model and electron density map (Fig. 7B and C, compare to Fig. 6B and C).

Outside the RBD regions, the trimeric RDR Env model is generally similar to the trimeric MLV Env model, including the intersubunit disulfide bond and hydrophobic SU anchor regions and SU helix αPD1 (Fig. 7F and G, compare to Fig. 4C and G). The only significant differences between the two models are in the membrane-proximal region. Whereas in the trimeric MLV Env model the αTM3 helices extend proximally parallel to the trimer axis and never contact each other, in the RDR Env model, the αTM3 helices converge and contact each other just before the MPER. This is associated with another significant difference between the models. Instead of a set of bulky SU-derived αPD5 helices separating the αTM3 helices, the RDR Env trimer model has the corresponding terminal SU regions in extended conformations with two additional β-strands, β13 and β14, in each subunit, allowing closer positioning of the αTM3 helices at the base of the trimer (Fig. 7B, C and F). The location of the last residue of SU and the first residue of TM in within a protomer and the configuration of the terminal SU regions crossing the trimer axis (Fig. 7B) implies that cleavage of Env by furin-like proteases and significant conformational changes must also occur before the final structure of the base of the trimer ectodomain is formed in RDR Env. Finally, a short αTM4 helix at the base of the trimer ectodomain parallel to the membrane plane forms an MPER with the side-chain of Phe-476 projecting into the membrane plane.

### Trimeric alpharetroviral Env models have composite structural features

The major differences between the Env of alpharetroviruses and gammaretroviruses are the absence of an independent RBD and an CXXC motif in the SU of alpharetroviruses. Instead, an amino-terminal Cys residue in the SU of alpharetroviruses forms a covalent disulfide bond with TM, similar to EBOV GP1 (Jeffers et al., 2002; Pike et al., 2011). Another similarity between alpharetroviral Env and filoviral SP not shared by the gammaretroviral Env is the presence of a disulfide loop in the fusion peptide in TM and GP2 (Delos and White, 2000; Lee et al., 2008). Modeling of trimeric alpharetroviral Env was performed to compare its structure to the two other major retroviral gamma-type trimeric proteins. Modeling was performed using the Env sequence of an ALV strain as described for the MLV Env trimer, retaining a linker between SU and TM during modeling.

Reliability scores for the trimeric ALV Env trimer model were generally lower than for the MLV and RDR Env models (Fig. 8A, C and D). In fact, only a small subset of ALV sequences tested yielded SU or trimeric Env models of reasonable quality. Despite the lower reliability scores, the *ab initio* trimeric ALV Env structural model provides sufficient information when interpreted in the context of the trimeric EBOV SP structure and MLV Env model.

**Figure 8.**
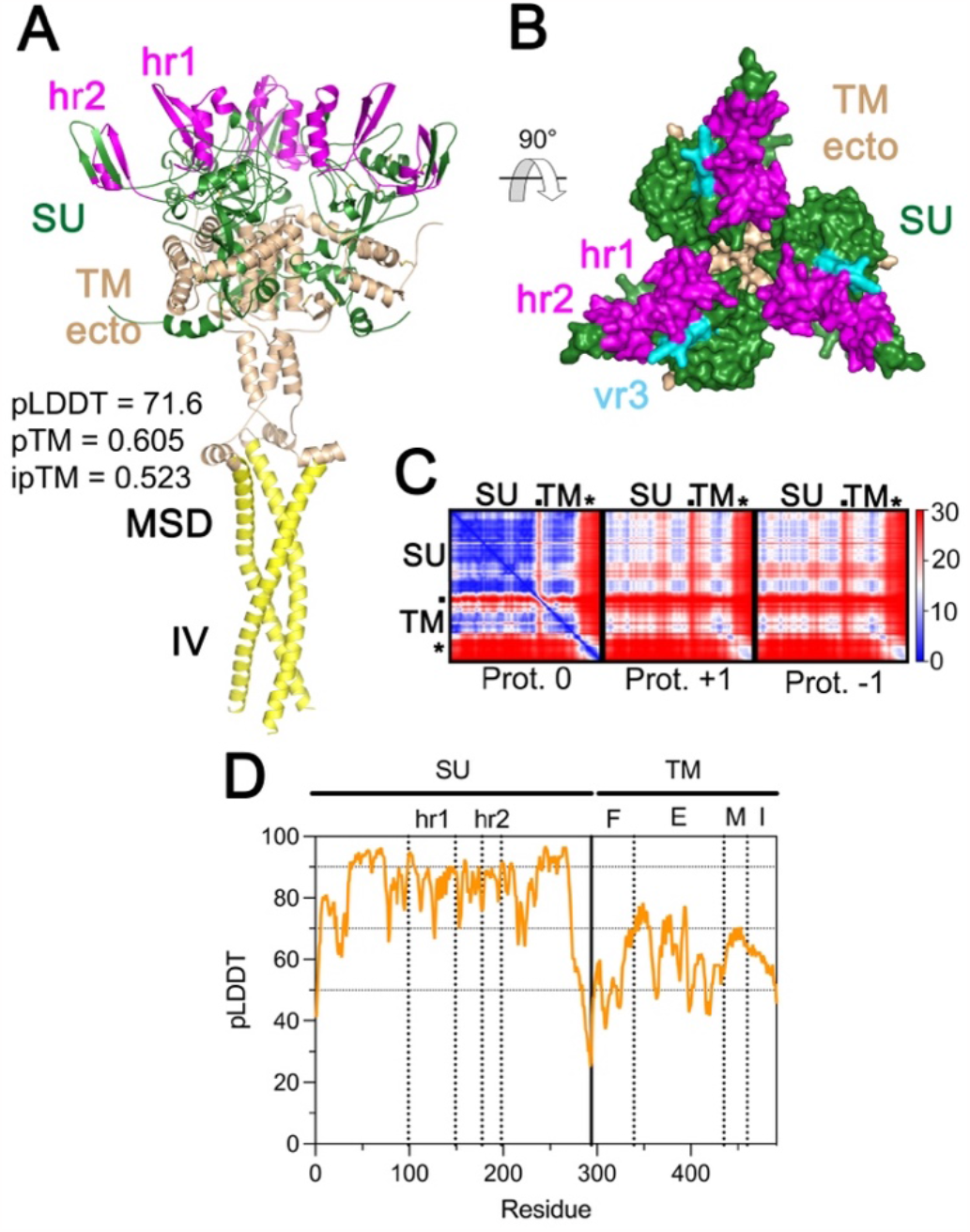
Trimeric ALV Env model. (A) Side view of the trimeric ALV Env model. SU is shown in green, with the host receptor interacting regions 1 and 2 (hr1 and hr2) of SU highlighted in magenta. The extracellular region of TM (TM ecto) is shown in wheat and the MSD and IV regions in yellow. AF2 quality scores for trimeric Env model are indicated in the panel. (B) Top view of the ALV Env trimer model shown in surface rendering. The extended variable region vr3 is shown in cyan. The (SG_3_)_3_ linker between SU and TM is omitted in panels A and B. (C) PAE scores of the trimeric ALV Env model. The leftmost panel indicates PAE scores within each protomer (“Protomer 0”), with the approximate location of the main Env regions shown on the left and top of the matrix. The dot indicates the location of the fusion peptide in the matrix and the asterisk the MSD. The middle and right panels show the PAE of “protomer 0” relative to the protomers in the counterclockwise (−1) and clockwise (+1) directions as observed from the top of the trimer. The PAE scale, in Å, is shown to the right of the matrices. The other effectively redundant matrices for each of the two other protomers in the trimer are not shown for simplicity. (D) Positional pLDDT scores of the ALV Env model. The boundaries or the major SU and TM regions are shown by vertical dotted lines. F, E, M, I indicate the fusion peptide, ectodomain, membrane-spanning domain and intraviral regions of TM. The threshold pLDDT scores of 50, 70 and 90 are shown by dotted lines.

The trimeric ALV Env model is similar to the trimeric MLV and RDR Env models without the RBD (Fig. 8A). In the ALV Env trimer model, two discontinuous loops corresponding to host-range regions 1 and 2 (hr1 and hr2) as well as variable region 3 (Holmen and Federspiel, 2000; Taplitz and Coffin, 1997) emanate from the PD distally to form a nearly contiguous surface at the distal end of Env in a position that would be available for receptor binding (Fig. 8A and B). The relative arrangement of α-helical regions in the ALV Env trimer model is more similar to the trimeric EBOV SP than to the MLV Env trimer model, especially in the αTM3 helices which, similar to the corresponding α-helices of the EBOV SP trimer, are located immediately around the vertical trimer axis and interact with each other through hydrophobic interactions (Fig. 3B and F, compare with Fig. 3A and E). The ALV Env trimer model also has short hydropathic membrane-proximal α-helix αTM4 homologues (Fig. 3B). These are positioned parallel to the envelope membrane and extend the hydrophobic surface of the membrane-spanning region as in the MLV and RDR Env trimer models, with large hydrophobic residues projecting towards the membrane, suggesting that this structure is a more general feature of gamma-type envelope proteins. The PD orientation in the ALV Env trimer model is the same as for the MLV Env trimer model and EBOV SP trimer structures (Fig. 3J). The fusion peptide forms a disulfide loop similar to the fusion peptide of EBOV GP2 on the surface of the trimer (Fig. 4E and F). The low pLDDT scores of that region may reflect flexibility of the fusion peptide in trimeric Env.

The ALV Env model has a short amino-terminal helix αPD1 positioned in a similar location and orientation in the trimer as the homologous MLV α-helix (Fig. 4F and G). The PD-proximal hydrophobic anchor around PD β-strands 2 and 11 is similar to the EBOV SP trimer structure and gammaretroviral Env models (Fig. 4B and F). The virion-proximal region of the hydrophobic anchor is similar in the ALV and the MLV and RDR Env trimer models, with the intersubunit disulfide bond located in a similar position in the alpha and gammaretroviral Env trimer models (Fig. 4B, C, F and G). In fact, the SU cysteine mediating the intersubunit disulfide bond in ALV Env (Pike et al., 2011) is located in a β-strand homologous to MLV PD strand β1 with the CWLC motif in MLV and RDR Env. Thus, the intersubunit disulfides are structurally homologous in the gammaretroviruses and ALV. However, the ALV Env model does not have a well-defined terminal PD β-strand 12 and the corresponding region in ALV Env adopts an extended conformation instead (Fig. 4F). The intersubunit disulfide is occluded in the virion-proximal hydrophobic anchor by an α-helix in the carboxy-terminal region of SU (Fig. 5B). Thus, the ALV Env trimer model has a structure intermediate between the gammaretroviral Env trimer models EBOV SP trimer structures, with an SU and intersubunit disulfide anchoring structure that is similar to the former and a fusion peptide and αTM3 configuration more similar to the latter. Instead of an unpaired Cys residue in a CXXC motif, the ALV Env model has a highly conserved unpaired SU Cys residue (Cys-34 in the model) that is critical for infectivity and is exposed only upon receptor binding (Smith and Cunningham, 2007) occluded in a mostly hydrophobic shell within the virion-proximal anchor, about 10 Å from the intersubunit disulfide bond (Fig. 5B).

## DISCUSSION

Here, AF2 multimer was used to understand the structure and evolution of trimeric retroviral Env. The quality of the trimeric Env models described here as assessed by the AF2 quality scores vary, high for the MLV and RDR Env models and lower for the ALV Env trimer model. These models can be used to understand the evolution and certain functional aspects of trimeric retroviral Env when considered together and in combination with experimental EBOV SP and HIV-1 Env trimer structures. The results show structural features that, in hindsight, are not particularly surprising and fit what might be expected by assuming certain key structural features to be highly conserved among gamma-type Env variants, including EBOV SP trimers, generally supporting the validity of the trimeric Env models. Furthermore, the structural models are compatible with all available structural and biochemical biological available for these proteins. The most surprising finding was the position and orientation of the RBD in MLV Env trimers, but also compatible with previous structural data. Thus, the models also reveal unexpected structural and functional features of retroviral Env.

The key model described here is the one for trimeric MLV Env, which has more extensive experimental data available to assess its validity. Although the trimeric MLV Env electron density maps were not used to inform modeling of the MLV Env trimer, the resulting model fits remarkably well in the best electron density maps. The model fitting results show that the head region of trimeric Env includes the C-domain rather than the RBD as previously proposed (Riedel et al., 2017; Wu et al., 2008). It should be noted that the previously proposed position of the RBD at the top of the trimer in electron density maps was not based on biochemical or structural data but rather on the expected position of a domain interacting with cell receptors. The trimeric RDR and ALV Env models also have the PD regions in a similar location in the distal region of Env. The location of the PD in the head region rather than in more membrane-proximal regions of Env as previously proposed is consistent with the location of the PD domain of EBOV GP1 in the trimeric SP structure. This would be expected, given the importance of this region in interacting with TM, which should be evolutionarily conserved across widely divergent gamma-type Env structures.

The trimeric Env models provide a detailed picture of the evolution of Env trimers in the retroviruses and filoviruses. First, as might be expected, the gamma-type Env share a common structural plan, including disposition of α-helical regions within pre-fusion TM/GP2, location of the intersubunit covalent disulfide bond in the trimer, configuration of the upper, PD-proximal hydrophobic SU anchor and PD orientation within the trimer. This conserved structural plan arises *de novo* in modeling of trimeric MLV, RDR and ALV Env, without templates from gamma-type Env structures, supporting these as shared features of gamma-type Env. However, there are clear distinctions between the different gamma-type Env models within the retroviruses and between these and the EBOV SP trimer. The retroviral gamma-type Env models have an additional SU helical region, αPD1, that replaces a small β-hairpin loop observed in a similar position in the EBOV SP. This α-helix, along with other slight rearrangements in TM, extend the hydrophobic anchor proximally towards the virus surface, forming the hydrophobic shell encasing the intersubunit covalent bond in the retroviruses. The ALV Env trimer seems to represent a structural intermediate between EBOV SP and MLV Env. The trimeric ALV Env has essentially the same overall structure as the EBOV SP trimer, including the fusion peptide largely within a disulfide loop, adding only a relatively short αPD1 helix in SU and minor changes in the SU carboxy-terminus that bury the intersubunit disulfide bond. The MLV and RDR Env trimer extends these modifications to add a cysteine in SU to form the CWLC motif within that structure. The distal anchoring structure is absent in HIV-1 Env, which instead has only a virion-proximal anchor in a similar position as the virion-proximal section of the anchor in MLV and RDR Env, although not completely homologous structurally, especially in the gp41/TM region. Of note, the apparent sequence of structural changes described here do not imply direct descent of SP/Env between these major viral groups.

The main objective in this study was to understand retroviral Env structural evolution. However, the models provide some surprising functional features of retroviral Env. Here, only the main functional consequences suggested by the models are briefly discussed.

The most important unexpected functional consequence that emerges from the models described here is the apparent functional RBD occlusion in trimeric MLV Env. The results of model fitting into trimeric MLV Env electron density maps indicate that the unusual tripod shape of the MLV Env trimer is given by the RBD, which can only fit in the outer regions of the tripod-shaped trimer when other trimer regions are fitted in the electron density maps, with the receptor-binding regions pointing away from the target cell. A lower-resolution electron density map of MLV Env bound to cell receptors in the context of target cell membranes show a chalice-like Env structure without the outer tripod legs, but instead with an extended density beyond the head region presumed to correspond to the RBD bound to the cell receptor (Riedel et al., 2017). This suggests a functional model in which the resting RBDs in trimeric Env sit retracted at the periphery of the trimer, shifting to an extended exposed configuration for receptor binding and to interact with the C-domain in the head region of the trimer to trigger fusion (Lavillette et al., 2001). In addition, type-C gammaretroviral Env can be functionally complemented by soluble RBD provided in *trans*, even when trimeric Env retains its own RBD moieties (Barnett et al., 2003, 2001; Barnett and Cunningham, 2001; Lavillette et al., 2001). The retracted RBD conformation would allow the soluble RBS moieties to directly interact with non-RBD Env regions, most likely the loop F region of the PD that is exposed on the head region of the trimer (Hötzel, 2022; Lavillette et al., 2001), to induce conformational changes in Env that trigger membrane fusion. The signals and structural mechanisms that would trigger the RBD of Env to move from the side to the apical end of Env to interact with cell receptors are not clear from the models or inspection of the lower-quality Env trimer models with RBDs extending distally at the top of the trimer but may involve the PRR (Lavillette et al., 1998). RBD retraction may participate in immune evasion by sequestration of receptor-binding sites away from neutralizing antibodies until the virus is close to the target cell, where antibodies may not be able to access critical epitopes. Of note, the cleavage site between SU and TM is buried by the RBD in the retracted state. This suggests that not only positioning of αPD5 close to the trimer axis but also the retracted RBD state may depend on Env cleavage by cellular proteases.

The trimeric RDR Env model provides the first view of key structural differences between this Env type and that of type-C retroviruses such as MLV. Although the SU region of the Env of these two retroviral groups is superficially similar, with an amino-terminal RBD and C-terminal C-domain-like separated by a CXXC motif, not only are the RBD moieties structurally distinct (Barnett et al., 2003, 2001; Hötzel, 2022) but the RBDs associate with the rest of the Env trimer differently in both groups and assume different positions in the trimer, a retracted configuration in the type-C Env and a more distally-exposed position in the RDR group. Thus, the structural models shown here point to major unexpected structural differences in the way the Env of these two gammaretroviral groups interact with host receptors. How the RBDs of these two gammaretroviral groups are related evolutionarily, by RBD modification or replacement, remains to be determined.

The results presented here show the value of novel deep-learning-based methods for structure prediction to advance the understanding of retroviral Env evolution and function, beyond what is possible by sequence analysis. Modeling of trimeric Env structures with AF2 multimer remains a challenge however, dependent in most cases on the identification of optimal starting sequences and sequence fragments and modeling strategies empirically, analogous to experimental approaches for structure determination. Some notable cases, such as Syn-1, have only yielded uninformative low-quality trimeric structural models resembling unfolded structures or TM conformations in the post-fusion state. In the case of Syn-1 the RDR Env structural model described here should provide a relatively close approximation for its trimeric structure and features including RBD position and orientation within the trimer. Modeling success with AF2 multimer may depend on the spectrum of sequence variants available in databases for a given retroviral group as well as selection of an optimal sequence within the group for modeling. The methods described here should inform strategies for modeling of other trimeric retroviral Env and syncytin structures. The structural models described here should also provide useful information to guide future experimental efforts to determine monomeric and trimeric Env structures, a major challenge in understanding the structure, function and evolution of these proteins.

## MATERIALS AND METHODS

### Trimeric MLV Env structural modeling

Structural modeling of trimeric polytropic MLV Env (GenBank XP_036020509) was performed with AF2 and AF2 multimer using the ColabFold server (Mirdita et al., 2022), accessible at https://colab.research.google.com/github/sokrypton/ColabFold/blob/main/AlphaFold2.ipynb with A100 graphics processing unit enabled. Selection of the MLV for Env modeling was based on preliminary monomeric and trimeric Env structural modeling using a diverse set of sequences of MLV Env strains, choosing the sequence that yielded the highest AF2 quality scores and lowest rate of modeling artifacts such as domain swaps. The segments used for modeling monomeric and trimeric Env, the corresponding templates and linker sequences between SU and TM, when used, are shown in Figure 1A. Monomeric Env modeling with standard AF2 used three recycles for each of the 5 modeling runs, without amber relaxation. The highest-scoring model (by pLDDT score) was selected as template for the next round trimeric Env structural modeling. Model regions omitted from templates were removed from PDB files using PyMol version 2.5.1 (Schrödinger LLC) prior to the next round of modeling. Trimeric Env modeling was performed using AF2 multimer v3. Two different trimeric Env modeling runs were performed in different dates to avoid re-use of the same random seed in different runs that result in identical models. The user-specified parameters used for modeling were the following: no amber relaxation; msa_mode: mmseqs2_uniref_env; model_type: alphafold2_multimer_v3; num_models: 5: num_recycles: null; recycle_early_stop_tolerance: null (default, 0.5 Å); rank_by: multimer; pair_mode: unpaired_paired; stop_at_score: 100.0; random_seed: 0; num_seeds: 1; use_dropout: false; use_cluster_profile: true. The second run to obtain models 3a and 3b used the same parameters except that it used 2 seeds for modeling (for a total of 10 final models). Ranked models were visually inspected to exclude models with TM resembling post-fusion states, if any, and chain-swapping artifacts between αTM1 and αTM2 or between αTM2 and the CX_6_CC motif loop. Modeling of trimeric EBOV SP was performed in a similar manner, to the trimeric Env modeling up to models 2a and 2b, using templates as described in the results section. Models were visualized with PyMol version 2.5.1.

### Modeling of trimeric RDR Env

Modeling of trimeric RDR Env was performed as described for trimeric MLV Env modeling but skipping the step modeling trimeric Env with a linker sequence between SU and TM. An endogenous *Rhinopithecus bieti* Env sequence (GenBank accession XP_017706409, residues 25 to 571) was selected for modeling following preliminary modeling runs with diverse exogenous and endogenous RDR Env sequences. The first round of monomeric Env modeling with AF2 used the sequence from residues 25 to 506 with a (G_3_S)_2_ linker between SU and TM residues 379 and 380. The second round to model trimeric Env with AF2 multimer used SU residues 25 to 379 and TM residues 380 to 571 separately and the top-scoring model from round 1 without low pLDDT regions and the linker sequence (residues 41 to 361 and 380 to 497) as template followed by amber-relaxation. Numbering of RDR Env in the final model descriptions sets residue 26 of XP_017706409 as residue 1.

### Modeling of trimeric ALV Env

Modeling of trimeric ALV Env was performed as described for MLV Env models 2a and 2b, followed by amber relaxation of the top-scoring model. An ALV subgroup J Env sequence (GenBank QGW52078, residues 69 to 559) was used for trimeric Env modeling based on preliminary SU modeling with a diverse set of ALV Env sequences, selecting the sequence yielding the highest AF2 quality scores. An (SG_3_)_3_ linker sequence was inserted between the SU and TM sequences for monomeric and trimeric Env modeling. Residue numbering in the final model starts in residue 69 of QGW52078 set as 1, excluding the linker sequence for numbering of TM regions.

### Assessment of structural model quality

The final, amber-relaxed, trimeric Env models were assessed for structural clashes and overall structural quality attributes using the Molprobity server (http://molprobity.biochem.duke.edu) (Davis et al., 2007).

### Electron-density model fitting

Fitting of trimeric MLV model 3a excluding the RBD and PRR regions to the electron-density map of Electron Microscopy Data Bank (EMDB) accession code EMD_3373 was done using ChimeraX 1.6.1 (Goddard et al., 2017). The trimeric Env model section was initially fitted manually followed by automated fitting based on local alignment between the model and the electron-density map. Fitting of MLV RBD (PDB 1AOL) to the electron-density map in different alternative poses was performed manually.

### Structural alignment database searches

PDB files of single Env protomers from the trimeric Env models were used to search for similar structures in the PDB and AFDB using the DALI (http://ekhidna2.biocenter.helsinki.fi/dali/) (Holm, 2020) and FoldSeek servers (https://search.foldseek.com/search) (Kempen et al., 2023).

## Data availability

The coordinate file of trimeric RDR, MLV and ALV Env models are available in the ModelArchive (https://www.modelarchive.org) with accession codes ma-x4xpz, ma-ni5co (https://modelarchive.org/doi/10.5452/ma-ni5co, using access code ihgKTMQg5X), and ma-xd0el (https://modelarchive.org/doi/10.5452/ma-xd0el, using access code jFdLCNalO3).

